# Discordance between biomarkers of iron status in plasma and myocardium across iron replete, iron deficiency, and iron replacement states

**DOI:** 10.1101/2024.11.10.622860

**Authors:** Kristina Tomkova, Katerina Cabolis, Maria Sajic, Marcin J Wozniak, Toby Richards, Gavin J Murphy

## Abstract

It is unclear whether plasma measures of iron status reflect iron status in tissues. This study compared the effects of iron deficiency or treatment with intravenous iron on plasma versus myocardial measures of iron metabolism and mitochondrial dysfunction in mice and humans. The human study included 31 people undergoing cardiac surgery. Twenty-two (71%) were iron deficient. The mouse study included 88 mice randomly allocated to iron replete diet group (Controls, n=29), an Iron Deficient diet (n=31) or an iron deficient diet followed by IV Iron administration (n=28). Outcomes included plasma iron and ferritin concentrations, and myocardial total iron, ferritin, IRP1 binding, and mitochondrial complex activity and DNA damage. In the human study, plasma ferritin positively correlated with myocardial ferritin (r^2^=0.155, p=0.012). People with iron deficiency had significantly lower myocardial ferritin levels than those without iron deficiency. There were no differences or significant correlations for other outcomes. In the mouse study, plasma ferritin negatively correlated with myocardial ferritin (r^2^=0.113, p=0.004), and iron deficiency reduced myocardial ferritin levels (p=0.018) as well as the binding affinity of myocardial IRP1 to the Ferritin heavy chain mRNA (p=0.033). Intravenous iron had divergent effects on plasma (increased) and myocardial (decreased) ferritin levels, and increased IRP1 binding affinity and mitochondrial respiratory complex activity (p=0.049). These finding suggest that measures of iron metabolism and iron deficiency in plasma do not consistently reflect myocardial iron status across iron replete, iron deficient, and iron replacement states.

## Introduction

In an apparent paradox, people with iron deficiency and anaemia who present for cardiac surgery are at increased risk of adverse outcomes [2], however these risks are not reduced by pre-surgery iron replacement [10]. Dysregulated iron metabolism is central characteristic of cardiovascular disease; for example, clinical progression of Heart Failure Reduced Ejection Fraction is associated with worsening iron restriction that ultimately progresses to iron deficiency, as determined by plasma biomarkers, in late-stage disease. Here again, iron replacement with intravenous iron also has not shown important clinical benefits in trials [7]. We hypothesised three possible explanations for these clinical observations. First, iron status measured in plasma may not reflect iron status in myocardium. For example, it has been established that almost all IV iron is rapidly taken up by the reticulo-endothelial system, where it is converted into plasma ferritin or metabolised as transferrin bound iron for utilisation by canonical cellular iron uptake pathways that are tightly regulated [3]. Whether these processes reverse the cellular effects of dysregulated iron metabolism in tissues is unclear, particularly in inflammatory states. Second, effective iron replacement in myocardium may not reverse cellular changes associated with iron deficiency such as mitochondrial dysfunction. Third, iron replacement may have unrecognised deleterious effects, especially IV ferric carboxy maltose, through oxidative mechanisms [16]. To test these hypotheses, this study compared the effects of iron replete status versus iron deficiency on simultaneous measures of iron metabolism in plasma and measures of iron metabolism, mitochondrial dysfunction, and DNA damage in myocardium from humans and mice, and evaluated the effects of intravenous iron replacement on these measures in iron deficient mice.

## Results

### Human study

A total of 40 people completed the VALCARD trial, of whom 31 provided blood samples for pre-surgery iron and ferritin analyses, and who constituted the analysis population. Of these, 25 provided adequately sized biopsies for assessment of myocardial iron and ferritin levels and mitochondrial complex assays, and 22/31 (71%) had plasma ferritin<30 ng/ml (**Figure 1A**).

**Figure 1.**
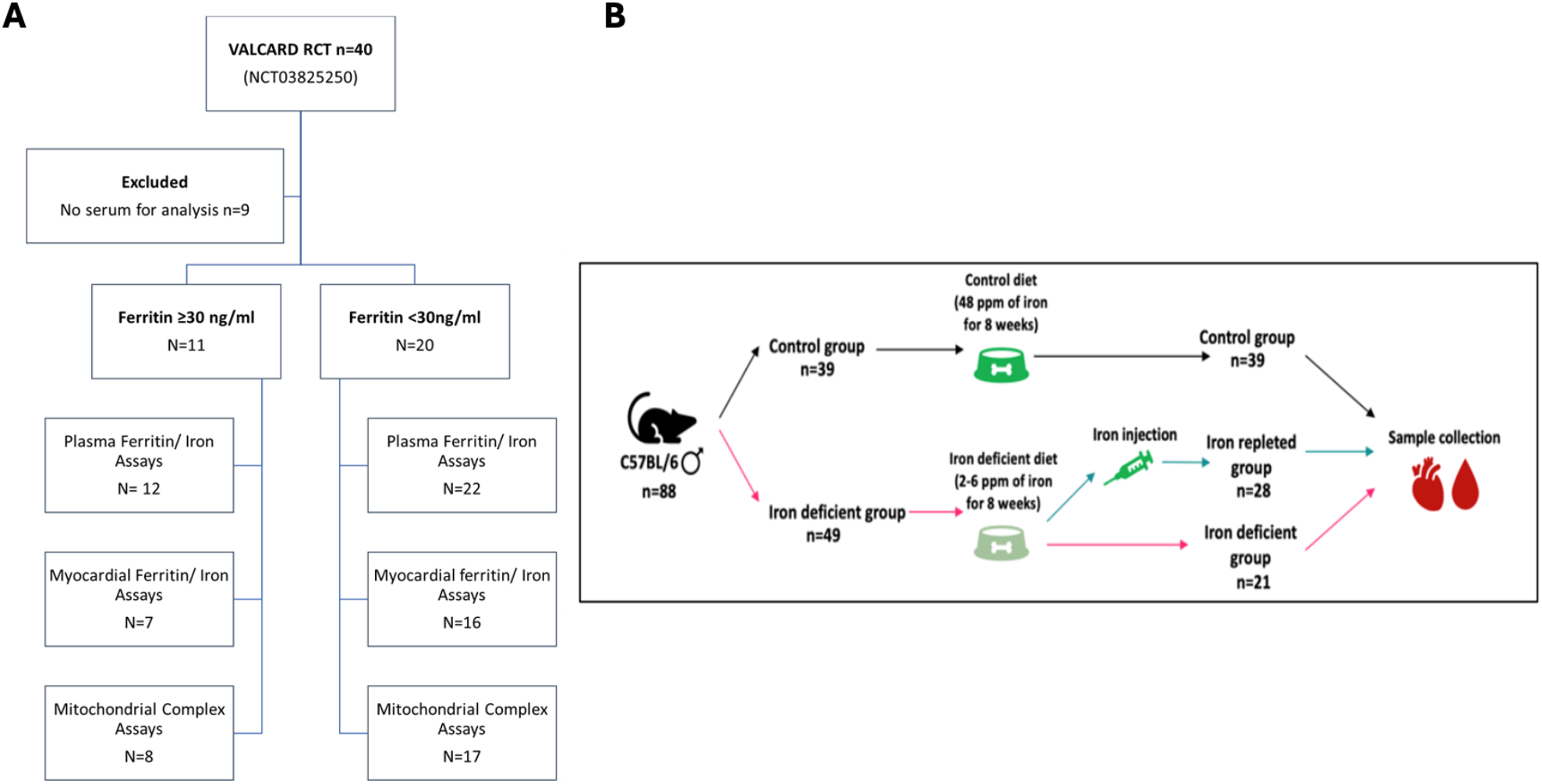
Study Schematics demonstrating the participant flow in the human study and study design in the mouse study. A) total of 40 patients were recruited for this study within the Valcard clinical trial. Out of these, 9 patients were excluded due to missing samples. The included patients were split into two groups based on their ferritin levels. On samples in both of these group three analyses were preformed, including the measurements for plasma iron and ferritin, myocardial iron and ferritin, and the function and expression of mitochondrial respiratory complexes. B) total number of 88 mice was used. These were split into two groups which received either control or iron deficient diet. After 8 weeks, the non-control group was further split into iron repleted group which received an intravenous iron injection, and iron deficient group which received an injection of saline solution. The sample sizes for each group are shown, as well as the diet details.

In bivariate analysis, there was a significant *positive* correlation between plasma and myocardial ferritin (r^2^=0.155, p=0.012). No other significant correlations between plasma and myocardial iron and ferritin levels, or with myocardial measures of mitochondrial complex activity or expression or mtDNA damage were demonstrated (**Figure 2**).

**Figure 2.**
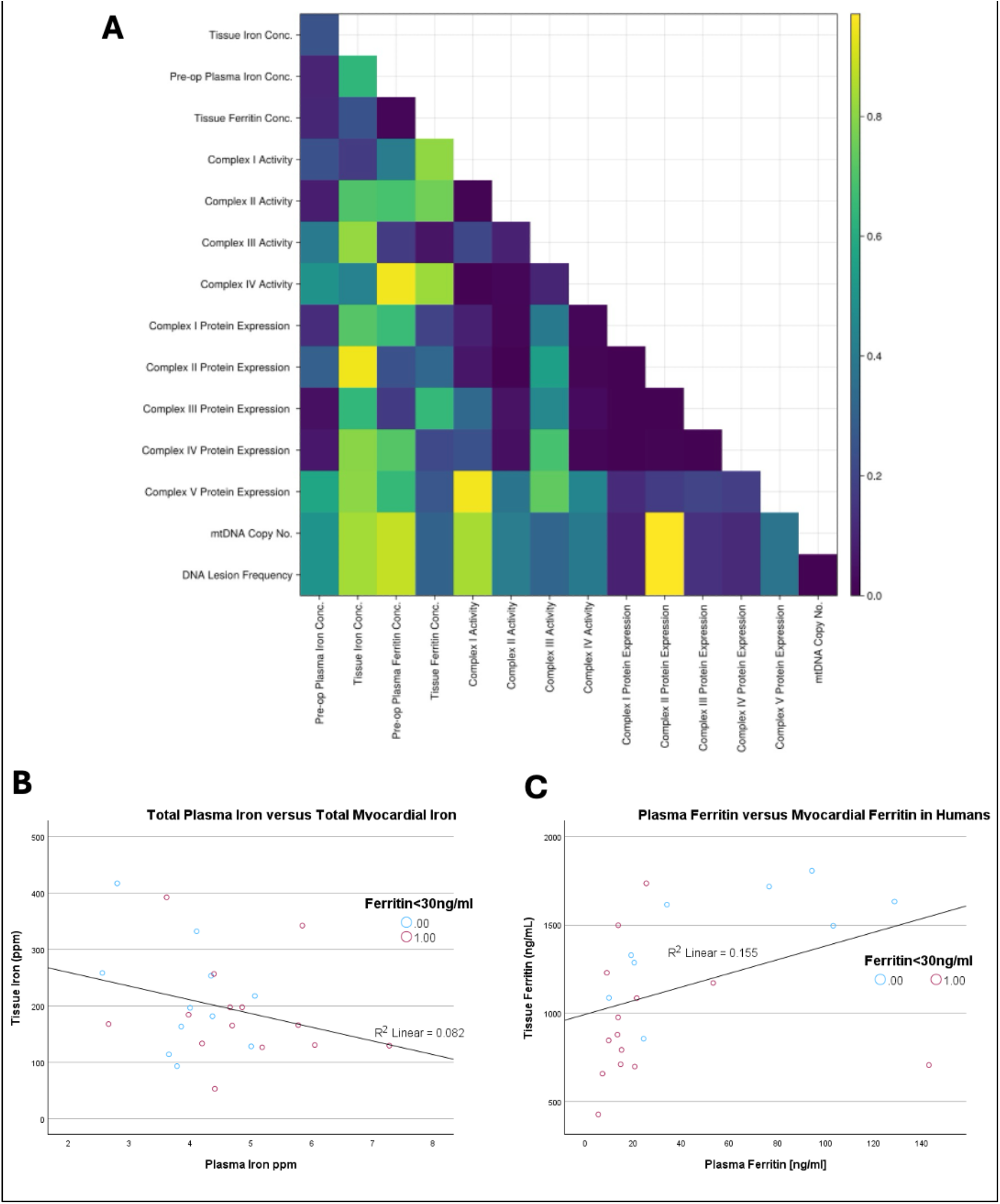
Results of the human bivariate analyses. A) Heatmap of Spearman correlation p Values from the bivariate analysis of the results of human study. B) Scatterplot showing Plasma Total Iron versus Myocardial Total Iron in Humans. C) Scatterplot showing Plasma Ferritin and Myocardial ferritin in Human

In subgroup analyses, participants with plasma ferritin< 30ng/ml had lower myocardial ferritin levels than participants with plasma ferritin ≥30ng/mL; MD 443.3ng/mL (107.8, 778.9), p=0.027 (**Table 1**). There were no differences between the groups for other outcomes.

**Table 1.** Measures of iron metabolism in blood and myocardium, and mitochondrial function in people with and without iron deficiency.

| Test<br>(Number of subjects) | Plasma Ferritin >30ng/mL<br>Median (IQR) | Plasma Ferritin <30ng/mL<br>Median (IQR) | Kruskal- Wallis<br>p Value | Control-Iron Deficient<br>Mean Difference (95%CI) |
| --- | --- | --- | --- | --- |
| Plasma Total Iron [ppm] n=30 | 4.0 (2.8, 5.0) | 4.3 (4.0, 5.2) | 0.250 | -0.582 (-1.40, 0.24) |
| Myocardial Total Iron (ppm) n=22 | 196.9 (128.7, 342.3) | 168.2 (129.8, 168.1) | 0.447 | 44.2 (-39.2, 127.5) |
| **Plasma Ferritin [ng/ml] n=31** | **76.5 (53.2, 128.6)** | **15.2 (13.5, 21.5)** | **<0.001** | **60.8 (38.4, 83.2)** |
| **Myocardial Ferritin [ng/ml] n=23** | **1614.2 (1170.4, 1716.4)** | **975.8 (845.6, 1327.9)** | **0.027** | **443.3 (107.8, 778.9)** |
| Complex I activity (nmol/min/kg) n=23 | 24.2 (22.6, 38.7) | 32.3 (19.3, 45.1) | 1.000 | -2.18 (-14.2, 9.8) |
| Complex II Activity (nmol/min/kg) n=23 | 43.6 (26.2, 52.3) | 34.1 (28.8, 47.1) | 0.720 | 4.8 (-11.2, 20.9) |
| Complex III Activity (nmol/min/kg) n=23 | 70.2 (33.7, 95.9) | 63.5 (27.0, 112.1) | 0.535 | 11.3 (-30.4, 52.9) |
| Complex IV Activity (nmol/min/kg) n=23 | 62.1 (56.8, 113.5) | 86.6 (70.3, 116.2) | 0.720 | -2.33 (-31.8, 27.1) |
| Expression Complex I (AU) n=23 | 0.64 (0.63, 0.80) | 0.68 (0.50, 0.89) | 0.671 | -0.60 (-0.32, 0.20) |
| Expression Complex II (AU) n=23 | 0.46 (0.45, 0.75) | 0.48 (0.43, 0.54) | 0.624 | 0.002 (-0.25, 0.25) |
| Expression Complex III (AU) n=23 | 0.77 (0.71, 0.87) | 0.96 (0.66, 1.06) | 0.103 | -0.19 (-0.43, 0.05) |
| Expression Complex IV (AU) n=23 | 0.99 (0.90, 1.05) | 0.90 (0.66, 1.11) | 0.588 | -0.022 (-0.33, 0.28) |
| Expression Complex V (AU) n=23 | 1.02 (0.83, 1.05) | 0.82 (0.53, 0.96) | 0.103 | 0.02 (-0.027, 0.33) |
| Normalised mtDNA (AU) n=25 | 18.8 (18.7, 23.1) | 21.5 (20.5, 24.0) | 0.588 | -0.90 (-3.70, 1.90) |
| mtDNA Lesions / 10 kb n=25 | 0.017 (-0.099, 0.142) | -0.017 (-0.0144, 0.035) | 0.588 | 0.52 (-1.11, 0.21) |

Raw data expressed as Median (Interquartile Range). P values derived from Kruskal-Wallis Test. For each inter-group comparison asymptotic significances (2-sided tests) are displayed with mean differences and 95% Confidence Intervals. The significance level is 0.050. AU, arbitrary units.

### Mouse Study

A total of 90 animals were used in the experiments. Two animals died prior to tissue harvest and were excluded from the analyses. The analysis cohort included 88 subjects. Of these 29 received an iron replete diet (200ppm) for 8 weeks (Controls), 31 received an iron deficient diet (95ppm) for 8 weeks with an intravenous saline infusion 48-72 hours prior to sacrifice (Iron Deficient), and 28 received an iron deficient diet (5ppm) for 8 weeks with IV iron administration (15mg/kg) 48-72 hours prior to sacrifice (IV Iron) (**Figure 1B**). By chance, baseline weight was higher in the control group compared to the two intervention groups, however this difference was maintained over the course of the study and demonstrated no interaction with treatment such that the differences in weights were similar at tissue harvest (**Supplementary Figure 1**).

In bivariate analysis there was a significant *negative* correlation between plasma and myocardial ferritin levels (r^2^=0.113, p=0.004). No other significant correlations were observed between plasma and myocardial iron and ferritin levels, or with myocardial measures of mitochondrial complex activity or expression or DNA damage (**Figure 3**).

**Figure 3.**
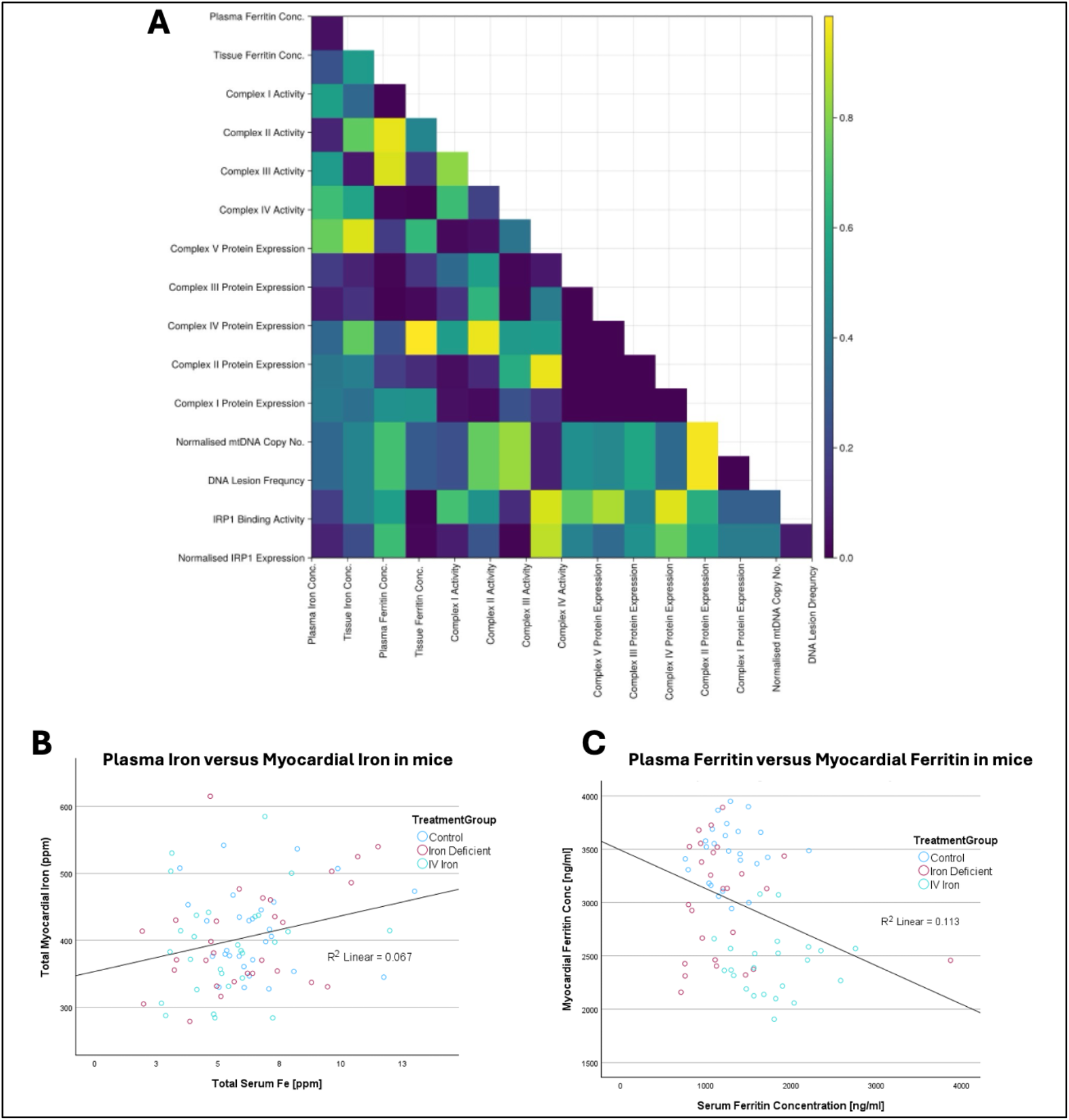
Results of the mouse bivariate analyses. A) Heatmap of Spearman correlation p Values from the bivariate analysis of the results of mouse study. B) Scatterplot showing Plasma Total Iron versus Myocardial Total Iron in mice. C) Scatterplot showing Plasma Ferritin and Myocardial Ferritin in mice

In the subgroup analyses, neither plasma total iron nor total myocardial iron concentrations were different between the three groups. Assessment of compartmentalised iron in the cytoplasm and the mitochondria in a sub-sample (n=39) also demonstrated no differences between the groups.

Plasma Ferritin concentrations were not different between Controls and Iron Deficient groups but were significantly higher in the IV Iron group versus the Control group; MD 566ng/ml, (257, 874), p<0.0001, and versus the Iron Deficient group; MD 633ng/mL, (329, 937), p<0.0001. Myocardial Ferritin concentrations normalised for total protein were significantly lower in the Iron Deficient group versus Controls; MD - 448ng/mL, (−184, -713), p=0.018. Iron replacement with Intravenous Iron resulted in a further drop in Myocardial Ferritin versus the Iron Deficient group; MD - 591ng/mL, (−321, -862), p=0.001.

The Iron Deficient diet had no significant effect on the activity of Mitochondrial Complexes I, II, III and IV versus Controls (**Table 2**). However, Intravenous Iron resulted in significantly higher activity of Complex III; MD 85.8 nmol/min/kg, (23.6, 1481), p=0.010, and Complex IV; MD 59.5 nmol/min/kg, (4.6, 1144), p=0.043, compared to Iron Deficient subjects. Intravenous Iron also resulted in significantly increased Complex III activity; MD 136.6 nmol/min/kg, (76.2, 1970), p<0.0001 versus Controls and significantly reduced Complex I activity versus Controls; MD -32.0 nmol/min/kg, (−6.2, -70.3), p=0.049. There was no difference between the groups with respect to levels of mitochondrial complex mRNA expression. There was no difference between the groups for mitochondrial DNA damage, a marker of oxidative stress, with adjustment for mitochondrial DNA concentrations (**Table 2**).

**Table 2.** Measures of iron metabolism in blood and myocardium, and mitochondrial function in mice stratified by subgroup.

| **Test**<br>(Number of subjects tested) | **Control**<br>Median (IQR) | **Iron Def Anaemia**<br>Median (IQR) | **IV Iron Therapy**<br>Median (IQR) | **Kruskal-Wallis**<br>p Value | **Control-Iron Deficient**<br>(Mean Difference, 95%CI, P value) | **Control-IV Iron**<br>(Mean Difference, 95%CI, P value) | **Iron Deficient- IV Iron**<br>(Mean Difference, 95%CI, P value) |
| --- | --- | --- | --- | --- | --- | --- | --- |
| Baseline Weight (g) n=88 | 20.3<br>(15.9, 21.8) | 18.2<br>(15.1, 19.6) | 18.9<br>(17.8, 20.8) | **0.038** | **1.50, (-0.22, 3.23), p=0.036** | 0.03, (-1.74, 1.79), p=1.000 | -1.48, (-3.21, 0.26), p=0.303 |
| Final Weight (g) n=88 | 27.90<br>(24.3, 30.5) | 25.15<br>(23.5, 27.1) | 25.50<br>(24.4, 28.4) | **0.011** | **2.44, (0.51, 4.37), p=0.020** | **2.15, (0.17, 4.13), p=0.037** | -0.29, (-2.24, 1.66), p=1.00 |
| Baseline Haemoglobin (g/L) n=87 | 142<br>(131, 151) | 142<br>(135, 159) | 143<br>(134-154) | 0.476 | -3.33, (-13.37, 6.71), p=0.672 | -1.77, (-12.16, 8.62), p=1.000 | 1.56, (-8.66, 11.79), p=1.000 |
| Final Haemoglobin (g/L) n=88 | 145.50 (137.25, 149.75) | 104.50<br>(92.5, 134.50) | 110.00 (102.50, 121.00) | **0.000** | **33.6, (21.8, 45.3), p<0.0001** | **35.3, (23.2, 47.3), p<0.0001** | 1.69, (-10.16, 13.55), p=1.000 |
| Plasma Total Iron [ppm] n=88 | 5.29<br>(4.36, 6.54) | 3.29<br>(2.60, 4.82) | 4.82<br>(3.06, 6.22) | 0.086 | 0.42, (-0.98, 1.82), p=0.517 | 1.21, (-0.23, 2.64), p=0.084 | 0.79, (-0.63, 2.20), p=1.000 |
| Myocardial Total Iron (ppm) n=86 | 432.0<br>(381.3, 482.1) | 398.3<br>(330.2, 429.5) | 393.1<br>(336.2, 470.6) | 0.489 | 8.6 (-38.1, 55.4), p=1.000 | 21.1, (-26.0, 68.3), p=0.723 | 12.5, (-34.7, 59.7), p=1.000 |
| Normalised Mitochondrial iron n=39 | 0.41<br>(0.35, 0.55) | 0.27<br>(0.14, 0.28) | 0.43<br>(0.42, 1.00) | 0.165 | -0.12, (-0.45, 0.21), p=1.000 | -0.17, (-0.56, 0.21), p=0.192 | -0.06, (-0.46, 0.35), p=0.380 |
| Normalised Cytoplasm iron n=39 | 1.03<br>(0.71, 1.53) | 1.12<br>(0.54, 2.01) | 1.29<br>(1.25, 1.32) | 0.537 | -0.18, (-0.74, 0.39), p=1.000 | 0.04, (-0.62, 0.70), p=0.798 | 0.22, (-0.48, 0.92), p=1.000 |
| Plasma Ferritin [ng/ml] n=88 | 1296.2<br>(1234.6, 1454.1) | 1198.0<br>(1034.4, 1561.6) | 1840.0<br>(1510.4, 2125.6) | **0.000** | 67.3, (-233.6, 368.1), p=0.530 | **-566, (-874, -257), p=0.000** | **-633, (-937, -329), p=0.000** |

| Test<br>(Number of<br>subjects tested) | Control<br>Median (IQR) | Iron Def Anaemia<br>Median (IQR) | IV Iron Therapy<br>Median (IQR) | Kruskal-Wallis<br>p Value | Control-Iron<br>Deficient<br>(Mean Difference,<br>95%CI, P value) | Control-IV Iron<br>(Mean Difference,<br>95%CI, P value) | Iron Deficient- IV<br>Iron<br>(Mean Difference,<br>95%CI, P value) |
| --- | --- | --- | --- | --- | --- | --- | --- |
| Myocardial<br>Ferritin [ng/ml]<br>n=74 | 3470.3<br>(3300.5, 3771.5) | 3131.0<br>(2566.0, 3664.5) | 2365.0<br>(2063.5, 2594.0) | **0.000** | **448, (184, 713),**<br>**p=0.018** | **1040, (767, 1313),**<br>**p=0.000** | **591, (321, 862),**<br>**p=0.001** |
| Normalised<br>IRP1 expression<br>n=72 | 1.58<br>(1.31, 2.33) | 1.43<br>(1.18, 1.85) | 1.34<br>(0.94, 1.68) | **0.015** | 1.12, (0.07, 2.18),<br>p=0.176 | **1.36, (0.31, 2.41),**<br>**p=0.012** | 0.23, (-0.77, 1.23),<br>p=0.925 |
| IRP1 Pull-down<br>Assay (IU)<br>n=85 | 0.00<br>(0.00, 598.38) | 0.00<br>(0.00, 0.00) | 0.00<br>(0.00, 189.80) | **0.009** | **622, (-2046, 3289),**<br>**p=0.018** | **2374, (-392, 5140),**<br>**p=0.033** | 1753, (-992, 4497),<br>p=1.000 |
| Complex I<br>activity<br>(nmol/min/kg)<br>n=71 | 200.0<br>(158.8, 207.3) | 170.8<br>(145.1, 226.7) | 193.6<br>(153.2, 255.6) | **0.029** | 5.4, (-31.5, 42.3),<br>p=1.000 | **32.0, (-6.2, 70.3),**<br>**p=0.049** | 26.6, (-12.3, 65.6),<br>p=0.071 |
| Complex II<br>Activity<br>(nmol/min/kg)<br>n=68 | 490.0<br>(379.6, 548.2) | 497.7<br>(407.5, 630.4) | 439.8<br>(371.3, 561.4) | 0.365 | -41.7, (-125.9, 42.5),<br>p=0.526 | -11.1, (-95.3, 73.1),<br>p=1.000 | 30.6, (-52.7, 113.8),<br>p=0.900 |
| Complex III<br>Activity<br>(nmol/min/kg)<br>n=74 | 138.0<br>(27.3, 177.5) | 113.5<br>(91.8, 266.0) | 182.7<br>(176.8, 303.8) | **0.000** | -50.8, (-111.8, 10.3),<br>p=0.145 | **-136.6, (-197.0, -**<br>**76.2), p=0.000** | **-85.8, (-148.1, -**<br>**23.6), p=0.010** |
| Complex IV<br>Activity<br>(nmol/min/kg)<br>n=77 | 207.0<br>(157.3, 273.3) | 171.0<br>(124.5, 190.6) | 248.1<br>(188.7, 336.8) | **0.046** | 36.5, (-18.9, 91.9),<br>p=0.342 | -23.0, (-78.5, 32.4),<br>p=1.000 | **-59.5, (-114.4, -4.6),**<br>**p=0.043** |
| Expression<br>Complex I (AU)<br>n=80 | 0.17 (0.10, 0.29) | 0.43 (0.06, 1.97) | 0.10 (0.06, 0.22) | 0.877 | -0.1, (-0.39, 0.19),<br>p=1.000 | -0.02, (-0.33, 0.28),<br>p=1.000 | 0.08, (-0.22, 0.38),<br>p=1.000 |
| Expression<br>Complex II (AU)<br>n=80 | 0.28<br>(0.24, 0.40) | 0.69<br>(0.30, 3.32) | 0.17<br>(0.13, 0.34) | 0.394 | -0.28, (-0.74, 0.18),<br>p=0.782 | -0.10, (-0.57, 0.37),<br>p=0.658 | 0.18, (-0.29, 0.65),<br>p=1.000 |
| Expression<br>Complex III<br>n=80 | 0.53<br>(0.34, 0.99) | 1.28<br>(0.46, 4.33) | 0.78<br>(0.57, 0.93) | **0.048** | -0.09, (-0.75, 0.57),<br>p=1.000 | -0.18, (-0.86, 0.50),<br>p=0.059 | -0.08, (-0.76, 0.59),<br>p=0.178 |
| Expression<br>Complex IV<br>(AU)<br>n=80 | 0.56<br>(0.42, 0.83) | 0.98<br>(0.47, 3.48) | 0.57<br>(0.35, 0.68) | 0.791 | 0.06, (-0.55, 0.68),<br>p=1.000 | 0.14, (-0.49, 0.76),<br>p=1.000 | 0.07, (-0.55, 0.70),<br>p=1.000 |
| Expression<br>Complex V (AU)<br>n=80 | 0.83<br>(0.25, 1.21) | 1.20<br>(0.48, 4.81) | 0.64<br>(0.43, 1.35) | 0.305 | -0.11, (-0.84, 0.63),<br>p=1.000 | 0.01, (-0.74, 0.77),<br>p=0.380 | 0.12, (-0.63, 0.87),<br>p=0.993 |
| Normalised<br>mtDNA (AU)<br>n=85 | 10.13<br>(8.79, 12.42) | 18.01<br>(15.03, 26.79) | 17.77<br>(15.80, 21.83) | 0.597 | -1.25, (-5.78, 3.28),<br>p=1.000 | -0.27, (-4.88, 4.35),<br>p=1.000 | 0.98, (-3.63, 5.60),<br>p=1.000 |
| mtDNA Lesions<br>/ 10 kb<br>n=85 | 0.36<br>(0.17, 0.48) | -0.17<br>(-0.51, -0.01) | -0.16<br>(-0.35, -0.05) | 0.597 | 0.14, (-0.14, 0.42),<br>p=1.000 | 0.01, (-0.28, 0.30),<br>p=1.000 | -0.13, (-0.42, 0.16),<br>p=1.000 |

Raw data expressed as Median (Interquartile Range). P values derived from Kruskal-Wallis Test. For each inter-group comparison asymptotic significances (2-sided tests) are displayed with mean differences and (95% Confidence Intervals). The significance level is 0.050 after the Bonferroni correction for multiple tests.
AU, arbitrary unit.

Iron Responsive Protein 1 (IRP1) concentrations normalised to tissue Tubulin concentration were lowest in the Intravenous Iron group versus Control; MD -1.36 AU, (−0.31, -2.41), p=0.012. Measurement of IRP1 binding capacity to the 5’ Iron Responsive Element (IRE) of Ferritin mRNA was significantly lower in the Iron Deficient group versus Controls MD -622 AU, (2046, -3289), p=0.018, indicative of low Iron Sulphur Cluster levels. This was reversed following IV iron administration; IV Iron-Iron Deficient MD IRP1 binding -2374 AU, (392, -5140), p=0.033 (**Table 2**).

## Discussion

In adults undergoing cardiac surgery, of whom 71% were iron deficient as defined by plasma ferritin <30ng/mL, there was a significant *positive* correlation between plasma and myocardial ferritin. No correlations were observed for other measures of iron metabolism. People with iron deficiency had significantly lower myocardial ferritin levels than those without iron deficiency, however there were no differences between the groups for other measures.

In the mouse study, there was a significant *negative* correlation between plasma and myocardial ferritin. No correlations were observed for other measures. There were no differences between the subgroups with respect to plasma or myocardial total iron. Dietary iron deficiency significantly reduced plasma ferritin compared to controls but there was no significant effect on plasma or myocardial iron or ferritin levels or on the expression or function of mitochondrial respiratory complexes. Iron deficient diets resulted in reduced binding affinity of myocardial IRP1 to the 5’ mRNA IRE for Ferritin mRNA. Intravenous iron administration in the setting of dietary iron deficiency had no effect on total plasma or total myocardial iron levels but had divergent effects on plasma (increased) and myocardial (decreased) ferritin. Intravenous iron increased availability of iron sulphur clusters (increased IRP1 binding affinity) and increased activity of mitochondrial complexes III and IV versus Iron Deficient mice.

This study has two significant findings. First, we demonstrated no differences in plasma total iron between iron deficient and non-iron deficient subjects whether mice or humans, or any correlation between plasma and myocardial total iron levels in mice and humans. In contrast, plasma and myocardial ferritin were significantly reduced in iron deficient subjects (both in mice and humans). The correlation between plasma and myocardial iron was positive in humans and negative in mice. This may be explained by the absence of an iron replacement group in the human study and the apparently paradoxical increase in plasma ferritin and simultaneous reduction in myocardial ferritin in mice who received intravenous iron. The reasons for this last finding are not clear from the current results. The Iron deficient diet did alter myocardial iron metabolism in mice; reductions in myocardial ferritin and low binding affinity of IRP1 to the 5’ mRNA IRE for Ferritin mRNA are evidence of reductions in the labile iron pool that is manifest as availability of iron sulphur clusters (ISC). IRP1 is a key regulator of cellular iron metabolism. In the presence of ISC excess, ISC bind to IRP1 increasing its affinity for the 5’ UTR IRE on Ferritin mRNA promoting translation, ferritin synthesis, and iron sequestration. Conversely, in iron deficiency, when ISC availability is low IRP no longer has affinity for ferritin mRNA but instead has affinity for the 3’ UTR for Transferrin Receptor 1 mRNA, promoting its translation and iron influx into cells [17]. Counterintuitively, we showed that increased IRP1 binding affinity for Ferritin heavy chain mRNA after IV iron was associated with a reduction in Myocardial Ferritin levels. To our knowledge, no similar study has reported myocardial ferritin in iron deficiency and 48-72hrs after IV iron administration for comparison. Here we hypothesise that other unmeasured suppressors of ferritin transcription such as oxidative stress [14] from acute influx of labile iron via L- and T-calcium channels as reported in a recent study [16] as a possible explanation. We did not detect mtDNA damage, a marker of mitochondrial oxidative stress, in our study however we did not measure oxidative stress across other cellular compartments, and multiple previous studies have demonstrated systemic and cellular oxidative stress post IV iron administration in mice [6].

Second, intravenous iron increased mitochondrial complex activity above that of iron replete controls. These changes were associated with indirect evidence of increased ISC availability as increased binding affinity for the 5’ mRNA IRE for Ferritin mRNA. Paradoxically, and for reasons that are unclear, the effects of IV iron were greatest on Complexes III and IV, whereas Complex I and Complex II and to a lesser extent Complex III are the most dependent on ISC for their function. Together, these observations support previous work demonstrating cardiac specific iron regulation that is not reflected by plasma markers of iron metabolism and identify the ongoing need for better research on the effect of iron deficiency and iron replacement on cellular iron metabolism in myocardial disease.

The study has three important limitations. First, the analysis did not directly measure the labile iron pool in mouse hearts which is a major determinant of cellular iron metabolism. We attempted to measure ISC levels in myocardium using an Electron Paramagnetic Resonance (EPR) Spectroscopy Technique, however this was not reproducible. Direct measurement of ISC has previously chiefly been reported only in cultured cells [5]. Our experience highlights the likely value of novel functional or imaging biomarkers of cellular iron metabolism and labile iron in myocardium for future research [8].

Second, the time points for tissue and blood analyses after iron administration used in this study (48-72 hours) are likely to have missed important acute changes in iron metabolism in blood and tissue. While we did not see any differences when comparing the 48- or 72-hours groups separately as a sensitivity analysis, Lakhal-Littleton and colleagues have shown that intravenous iron results in acute increases in myocardial iron concentrations within one hour of administration in parallel with acute rises in plasma and myocardial total iron [16]. Myocardial iron homeostatic mechanisms may therefore have normalised total cellular iron levels by 48-72 hours. In the Lakhal-Littleton study total myocardial iron concentrations were persistently higher at up to 42 days following intravenous iron versus iron deficient controls, whereas they were no different between groups at 48-72 hours in the current analyses. This difference may have been attributable to the slightly different design of the former study, where IV iron was administered at the start of the iron deficient diet, allowing myocardial uptake of RE delivered iron in the absence of dietary iron over the 42-day course of the study.

The third and major limitation is that dysregulated iron metabolism and mitochondrial dysfunction in myocardial diseases such as heart failure occur in the setting of inflammation and biological ageing [7]. The expression of Ferritin, and the major iron regulator hepcidin are increased by a number of inflammatory pathways including IL6-JAK2-STAT3 [15]. Pre-surgery IL6 levels in the human participants were low, and there was little evidence of systemic inflammation manifest as high ferritin in the absence of other factors, in the mouse study. Therefore, our findings may not be generalisable to the spectrum of dysregulated iron metabolism across myocardial disease.

Furthermore, there were important differences in the plasma ferritin levels, and in myocardial ferritin levels between mice and humans. Ferritin plasma levels in humans were 5-150 ng/ml whereas in mice the range was 700-4000 ng/ml, and atrial tissue ferritin was 350-1800 ng/ml in humans while myocardial tissue ferritin ranged between 1500-4000 ng/ml in mice. The former may be explained by multiple factors including age and comorbidity of the study subjects. In the latter case atrial iron levels in humans were compared to ventricular iron levels in mice which introduces further potential for heterogeneity. In mitigation, the associations between plasma and tissue iron indices were similar for both species. This represents a strength of the study despite differences in absolute concentration between species.

In the current study, plasma and myocardial ferritin levels were positively correlated in humans and negatively correlated in mice. Otherwise, changes in myocardial specific regulation of iron metabolism across iron replete, deficiency, and replacement states were not demonstrated by changes in plasma biomarkers of iron metabolism. Together, these results provide insights into the negative results of trials of iron replacement in people with cardiovascular disease and iron deficiency as defined by plasma biomarkers. The results support the wider use of novel tissue imaging and biomarker measures of tissue labile iron and molecular analyses of cellular responses to iron administration in human studies.

## Methods

### Human Study

Human data was obtained from participants in the VALCARD trial (NCT03825250), a dose finding randomised controlled trial of sodium valproate pre-cardiac surgery. All participants had provided informed consent for all analyses presented in this study; ethical approval was obtained prospectively from the East Midlands Research Ethics Committee (18/EM/0188). This secondary observational analysis is reported as per the Strengthening the Reporting of Observational studies in Epidemiology (STROBE) statement [18]. The STROBE checklist is included in the supplementary material. The study adhered to the principles outlined in the Declaration of Helsinki.

Adults undergoing coronary artery bypass grafting without pre-existing organ failure or a contraindication to sodium valproate were eligible to take part. Participants were allocated to receive Valproate at a dose of 15-25mg/kg for between 2 and 6 weeks. Sodium valproate had no effect on any if the outcome measures reported in this analysis. The primary analysis of the VALCARD trial will be reported separately. For the current analysis iron deficiency was defined as a plasma ferritin <30ng/ml. Blood samples were collected at anaesthetic induction, spun and plasma samples were frozen and stored at -80C. In addition, right atrial biopsies were collected in a standardized manner from the right auriculum before cannulation for cardiopulmonary bypass and were immediately snap-frozen in liquid nitrogen for later analysis. Only participants who had sufficient plasma and myocardial biopsy volumes after completion of the primary VALCARD analyses were included in the present study.

### Mouse Study

The mouse samples for this study were obtained from the University College London (Charles River, Harlow, UK and P-Block, UK). All animal handling and experiments were performed in accordance with the UK Home Office Animals (Scientific Procedures) Act 1986 and all experiments were approved by the institutional ethics committee at UCL and followed the ARRIVE guidelines (attached in the supplement).

A total of 90 male C57BL/6 mice were used. The sample size was decided based on past literature and pilot experiments. The experimental unit was considered to be a single animal. No exclusion/inclusion criteria were specified. The mice were weaned at 4 weeks. After arrival to the animal housing unit, the mice were allowed to acclimatise to the environment for a week prior to the beginning of the experiment and were culled after 8-9 weeks following the Code of Practice for the Humane Killing of Animals under Schedule 1. Mice were randomly allocated into cages in groups of five with access to food and water ad libitum, with a 12-hour light/dark cycle and at a consistent temperature of 21 ±2°C. The cages were randomly assigned to either control or iron deficient (ID) diet, and the specified diets were started from 5 weeks of age for 3-4 weeks. Mice in ID group (n=49) received an iron depleted diet containing 2-6 ppm of iron (Teklad Custom Diet, Envigo, Indianapolis, Indiana, USA). The mice in the control group (n=39) received identical diet supplemented with 48 ppm of iron (Teklad Custom Diet, Envigo, USA). From the ID group, 28 mice were randomly selected to receive iron repletion intervention to form the iron repleted group (IR). Repleted mice were injected intravenously with an emulsion of 15 mg/kg of ferric carboxymaltose (Ferinject, Vifor France, Paris, France) 24-72 hours before termination. Remaining mice were given a single intravenous injection of saline (Reliwash, Reliance Medical, UK) 48-72 hours before termination. To monitor the iron deficiency, haemoglobin (Hb) levels were assessed weekly by making a small incision to the tail vein using a 21G needle and collecting a drop of blood using specialised microcuvettes (HemoCue® Hb 201 Microcuvettes, 111715, Hemocue Radiometer, UK) and a point-of-care Hb analyser (HemoCue® 201+, 121717, Hemocue Radiometer, UK).

After termination, the mice were dissected, and blood and heart tissues were collected and stored as per human tissue. All personnel involved in sample processing and data analysis were blinded to which dietary group the animals belonged to.

### Measurements and outcomes

#### Total iron concentration in heart tissue and plasma

Total elemental iron concentrations in heart tissue and plasma in both mouse and human samples were measured using Inductively Coupled Plasma Mass Spectrometry (ICP MS), as previously described [11, 13]. The samples were washed thoroughly in PBS and processed using a microwave digest in nitric acid and hydrogen peroxide and the iron concentration was measured using the the iCAPq-c Quadrupole ICP MA machine (Thermo Scientific, USA) with the spiking solution containing 1 ppm iridium (PLIR3-2Y, Spex CertiPrep, USA) and 1 ppm of rhodium (7440-16-6, Sigma-Aldrich, USA). As the standard, the Multi-element Solution 2A (QX146627, Spex CertiPrep, USA) was used. Based on the standard, the total amount of iron was calculated, and the results were normalised to the dry tissue weight.

#### Ferritin concentration in heart tissue and plasma

Ferritin heavy chain concentration was measured using ELISA assays (Ferritin Mouse ELISA kit - KA4821, Mouse Ferritin FE ELISA kit -MBS261944, or Ferritin Human ab108837, Abcam, UK) following the manufacturer’s instructions. The standard curves and sample concentrations were calculated from the absorbance data using the MyAssays online software, using a Four Parameter Logistic (4PL) curve fit [4]. The results were normalised to the protein concentration of the homogenates obtained in a Bradford assay (5000006, Bio-Rad, UK).

#### Mitochondrial (mt) DNA copy number

mtDNA copy number was measured using quantitative real-time PCR. The protocol for this experiment protocol was adjusted from published literature. Genomic DNA (gDNA) was extracted from 15 mg (dry weight) of cardiac tissue using the DNeasy Blood & Tissue DNA extraction kit (69504, Qiagen, UK). The DNA concentration was measured using the PicoGreen assay (P7589, Invitrogen, USA). The DNA copy number was estimated using a standard curve produced from dilutions of plasmids of a known copy number. The master mix was supplied by Fisher Scientific (A25742) and the primers used for this experiment were as follows:

##### Mouse

*mtND1 gene*

Forward primer: TCCTAACACTCCTCGTCCCT

Reverse primer: ATGGCGTCTGCAAATGGTTG

*POLB gene*

Forward primer: CGTGCGGTTTGAGCTTTTGA

Reverse primer: AAATGGGGAAGCCGGTAAGG

##### Human

*RNR gene*

Forward primer: GGTTTGTTAAGATGGCAG

Reverse primer: GGAATTGAACCTCTGACTG

*B2M gene*

Forward primer: TGCTGTCTCCATGTTTGATGTATCT

Reverse primer: TCTCTGCTCCCCACCTCTAAGT

#### mtDNA damage

Mitochondrial DNA damage was measured using the Long-amplicon quantitative PCR (LA-QPCR) method, following the protocol previously published in literature. The LA-PCR master mix was supplied by the New England Biolab (M0287S) and the primer sequences were as follows:

Forward primer: GCCAGCCTGACCCATAGCCATAATAT

Reverse primer: GAGAGATTTTATGGGTGTAATGCGG

The DNA concentrations of technical replicates were averaged and normalised to their respective copy number. The results were expressed at the amplification relative to the control group.

#### Protein expression and enzymatic activities of myocardial mitochondrial respiratory complexes

To measure the enzymatic actives of mitochondrial respiratory complexes, a previously published protocol was adapted [12]. For this, a colorimetric assay was performed where the rate of colour change was directly proportional to the rate of substrate utilisation by the mitochondrial complexes. Complex specific substrates and inhibitors were used to determine the enzymatic activities of complexes I, II, III, and IV in tissue homogenate samples. The protein expression of these complexes as well as complex V was estimated using western blotting.

#### IRP1 binding activity in mouse hearts

The binding activity of the iron regulatory protein 1 (IRP1) protein to the iron responsive element (IRE) motif was assessed in a pull-down assay as described in previously published literature [9]. Biotinylated probes were produced using invitro transcription. The pull-down assay was performed on protein extract from cardiac tissue homogenates using streptavidin-conjugated magnetic beads. The results were visualised using western blotting and quantified using the ImageJ software by measuring the intensity of the IRP1 bands. The protein expression data was normalised to the total amount of protein loaded in the wells using the Ponceau S Staining Solution (A40000279, Fisher Scientific, USA)[1]. The DNS sequences of the oligonucleotides used for the RNA probes are as follows:

##### IRE probe

Forward primer: TTAAATTATGCTGAGTGATATCCCCAGAGTCGCCGCGGTTTCCTGCTTCAACAG

Reverse Primer: TGCTTGAACGGAACCCGGTGCTCGACCCCT

##### GAPDH probe

Forward Primer: TTAAATTATGCTGAGTGATATCCCGGATCCCCTGCTGGGAGGGGGCAGGG

Reverse Primer: GACCTGTTCCCACCGTGTGC

##### T7 promoter

AATTTAATACGACTCACTATAGG

### Statistical analyses

The data obtained in this study was analysed using SPSS 23.0 software (IBM, San Diego, California, USA). Raw data were presented as Median (Interquartile Ranges). Spearman correlation coefficients were estimated between blood and myocardial measures of iron metabolism and mitochondrial function. Subgroup analyses considered VALCARD participants with pre-surgery plasma ferritin <30ng/mL versus ≥30ng/mL, or individual treatment groups in the mouse study. To determine the significance of the differences between the examined subgroups, one-way Kruskal-Wallis was used. Intergroup treatment effects were expressed as Mean Difference (MD) and (95% Confidence Intervals). Pairwise subgroup comparisons were considered significant if the two-sided p value <0.05, with Bonferroni adjustment where required.

## Acknowledgements

The authors are grateful to funding bodies enabling the performance of this research as well as the research and administrative staff supporting this work.

The authors are also grateful to Prof Samira Lakhal-Littleton for her helpful comments on the manuscript.

## Details of authors’ contributions

KT performed all the laboratory analyses and co-wrote the manuscript.

TR conceived and designed the study.

KC and MS designed and performed the animal experiments.

MJW designed and supervised the laboratory analyses.

GJM conceived and designed the analyses, co-wrote the manuscript, and is the senior author.

All of the authors have reviewed and approved the final manuscript.

## Data availability statement

All data discussed in this study is presented in the form of tables and figures within the study or in the supplementary material. Further data can be provided by the corresponding author, upon reasonable request.

## Declaration of interests

GJM has worked as a Consultant for Pharmacosmos.

All other co-authors declare No Conflicts of Interest.

## Funding

The study was supported by British Heart Foundation Grants RG/17/9/32812, CH/12/1/29419, e and the Leicester NIHR Biomedical Research Centre (Cardiovascular)

## Notes

### Competing Interest Statement

G.J. Murphy has worked as a Consultant for Pharmacosmos.
All other co-authors declare No Conflicts of Interest.

