## supplementary material for "Discordance between biomarkers of iron status in plasma and myocardium across iron replete, iron deficiency, and iron replacement states"

**Supplementary Figure 1: Weekly weights in test subjects in the mouse study stratified by group.**

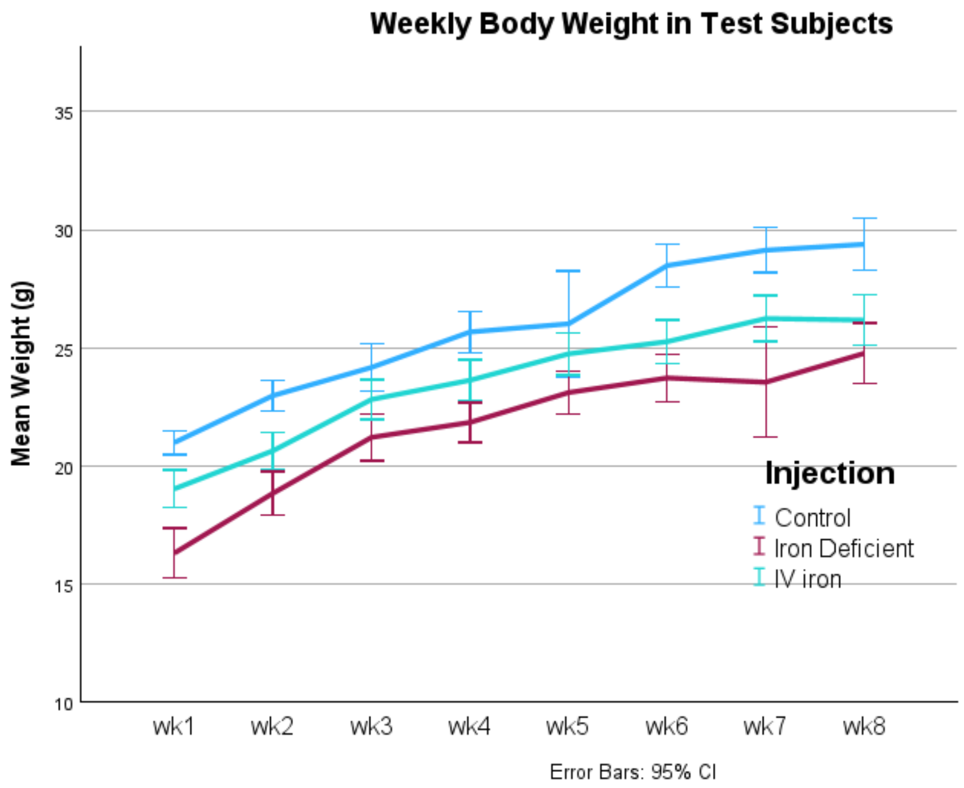

**Supplementary Table 1: Pre-surgery characteristics of the analysis cohort**

| | **All Participants** n=31 | **Plasma Ferritin >30ng/mL** n=11 | **Plasma Ferritin <30ng/mL** n=20 | p Value |
|---|---|---|---|---|
| Age | 66 (59.5, 73.5) | 69 (60, 75) | 65 (58.5, 71.25 | 0.317 |
| Male | 29 (93.5%) | 11 (100%) | 18 (90%) | 0.527 |
| White ethnicity | 28 (90%) | 9 (82%) | 19 (95%) | 0.303 |
| Body Mass Index | 28.0 (26.5, 29.7) | 27.6 (25, 28) | 28.3 (27, 30.1) | 0.212 |
| estimated Glomerular Filtration Rate (ml/min/1.73m$^2$) | 95 (77.2, 104.2) | 87 (75.8, 103.7) | 96.1 (82, 104.7) | 0.497 |
| Interleukin-6 (pg/mL) | 0.75 (0.40, 1.56) | 0.98 (0.52, 1.76) | 0.63 (0.46, 1.59) | 0.563 |
| New York Heart Association Class III/IV | 5 (16%) | 0 (0%) | 5 (25%) | 0.175 |
| Ejection Fraction <50% | 6 (19%) | 3 (27%) | 2 (15%) | 0.638 |
| Haemoglobin Concentration (g/L) | | 103 (96, 119) | 111 (98, 137) | 0.820 |
| Diabetes n=31 | 9 (29%) | 4 (36%) | 5 (25%) | 0.683 |
| COPD/Asthma | 1 (3%) | 0 | 1 (5%) | 1.000 |
| Neurological deficit | 1 (3%) | 0 | 1 (5%) | 1.000 |
| Frail | 6 (19%) | 3 (27%) | 3 (15%) | 0.638 |
| Operation type | | | | 0.542 |
| Isolated CABG | 16 (52%) | 6 (55%) | 10 (50%) | |
| Isolated valve | 12 (39%) | 3 (27%) | 9 (45%) | |
| CABG+Other | 3 (9%) | 2 (18%) | 1 (5%) | |

Continuous variables expressed as Median (Interquartile Range) P Value from Mann Whitney test. For ordinal data P values derived from Fisher's Exact test.
CABG, Coronary artery bypass grafts, COPD, Chronic Obstructive Airways Disease

### Supplementary Table 2: Correlation coefficients and p values of the analysis of the human study.

| | | Pre-op Plasma Iron Conc. | Tissue Iron Conc. | Pre-op Plasma Ferritin Conc. | Tissue Ferritin Conc. | Hb | Complex I Activity | Complex II Activity | Complex III Activity | Complex IV Activity | Complex I Expression | Complex II Expression | Complex III Expression | Complex IV Expression | Complex V Expression | mtDNA Copy No. |
|---|---|---|---|---|---|---|---|---|---|---|---|---|---|---|---|---|
| Tissue Iron Conc. | Correlation Coefficient | -0.24 | | | | | | | | | | | | | | |
| | Sig. (2-tailed) | 0.248 | | | | | | | | | | | | | | |
| | N | 25 | | | | | | | | | | | | | | |
| Pre-op Plasma Ferritin Conc. | Correlation Coefficient | -0.314 | -0.107 | | | | | | | | | | | | | |
| | Sig. (2-tailed) | 0.091 | 0.636 | | | | | | | | | | | | | |
| | N | 30 | 22 | | | | | | | | | | | | | |
| Tissue Ferritin Conc. | Correlation Coefficient | -0.33 | 0.26 | .516* | | | | | | | | | | | | |
| | Sig. (2-tailed) | 0.1 | 0.242 | 0.012 | | | | | | | | | | | | |
| | N | 26 | 22 | 23 | | | | | | | | | | | | |
| Complex I Activity | Correlation Coefficient | 0.239 | -0.314 | 0.182 | 0.05 | -0.261 | | | | | | | | | | |
| | Sig. (2-tailed) | 0.24 | 0.165 | 0.406 | 0.812 | 0.199 | | | | | | | | | | |
| | N | 26 | 21 | 23 | 25 | 26 | | | | | | | | | | |
| Complex II Activity | Correlation Coefficient | 0.358 | -0.088 | 0.088 | -0.064 | 0.09 | .579** | | | | | | | | | |
| | Sig. (2-tailed) | 0.073 | 0.703 | 0.69 | 0.762 | 0.662 | 0.002 | | | | | | | | | |
| | N | 26 | 21 | 23 | 25 | 26 | 26 | | | | | | | | | |
| Complex III Activity | Correlation Coefficient | -0.173 | -0.053 | 0.299 | .398* | -0.321 | -0.258 | -0.342 | | | | | | | | |
| | Sig. (2-tailed) | 0.397 | 0.819 | 0.165 | 0.049 | 0.11 | 0.203 | 0.087 | | | | | | | | |
| | N | 26 | 21 | 23 | 25 | 26 | 26 | 26 | | | | | | | | |
| Complex IV Activity | Correlation Coefficient | 0.141 | -0.182 | -0.013 | 0.045 | -0.153 | .657** | .476* | -0.334 | | | | | | | |
| | Sig. (2-tailed) | 0.494 | 0.43 | 0.954 | 0.829 | 0.454 | <.001 | 0.014 | 0.095 | | | | | | | |
| | N | 26 | 21 | 23 | 25 | 26 | 26 | 26 | 26 | | | | | | | |

| | | | | | | | | | | | | | | | | |
|---|---|---|---|---|---|---|---|---|---|---|---|---|---|---|---|---|
| Complex I Expression | Correlation Coefficient | 0.312 | -0.086 | -0.097 | 0.27 | 0.049 | 0.345 | .499** | 0.175 | .486* | | | | | | |
| | Sig. (2-tailed) | 0.121 | 0.712 | 0.66 | 0.192 | 0.81 | 0.084 | 0.009 | 0.391 | 0.012 | | | | | | |
| | N | 26 | 21 | 23 | 25 | 26 | 26 | 26 | 26 | 26 | | | | | | |
| Complex II Expression | Correlation Coefficient | 0.214 | 0.014 | 0.256 | 0.208 | -0.064 | 0.379 | .625** | -0.125 | .537** | .608** | | | | | |
| | Sig. (2-tailed) | 0.293 | 0.951 | 0.239 | 0.317 | 0.755 | 0.056 | <.001 | 0.544 | 0.005 | <.001 | | | | | |
| | N | 26 | 21 | 23 | 25 | 26 | 26 | 26 | 26 | 26 | 26 | | | | | |
| Complex III Expression | Correlation Coefficient | .406* | -0.108 | -0.3 | 0.097 | 0.029 | 0.201 | .398* | 0.154 | .436* | .871** | .649** | | | | |
| | Sig. (2-tailed) | 0.04 | 0.642 | 0.164 | 0.645 | 0.889 | 0.326 | 0.044 | 0.452 | 0.026 | <.001 | <.001 | | | | |
| | N | 26 | 21 | 23 | 25 | 26 | 26 | 26 | 26 | 26 | 26 | 26 | | | | |
| Complex IV Expression | Correlation Coefficient | 0.373 | -0.058 | -0.08 | 0.259 | 0.178 | 0.24 | .412* | 0.083 | .493* | .807** | .593** | .841** | | | |
| | Sig. (2-tailed) | 0.061 | 0.801 | 0.717 | 0.211 | 0.386 | 0.238 | 0.037 | 0.687 | 0.011 | <.001 | 0.001 | <.001 | | | |
| | N | 26 | 21 | 23 | 25 | 26 | 26 | 26 | 26 | 26 | 26 | 26 | 26 | | | |
| Complex V Expression | Correlation Coefficient | 0.115 | 0.057 | 0.101 | 0.235 | 0.041 | -0.011 | 0.183 | -0.07 | 0.156 | 0.315 | 0.277 | 0.262 | 0.275 | | |
| | Sig. (2-tailed) | 0.575 | 0.806 | 0.647 | 0.259 | 0.844 | 0.956 | 0.371 | 0.734 | 0.446 | 0.117 | 0.171 | 0.196 | 0.175 | | |
| | N | 26 | 21 | 23 | 25 | 26 | 26 | 26 | 26 | 26 | 26 | 26 | 26 | 26 | | |
| mtDNA Copy No. | Correlation Coefficient | 0.13 | -0.044 | 0.031 | -0.219 | -0.128 | 0.042 | 0.177 | -0.216 | -0.183 | -0.357 | -0.008 | -0.317 | -0.333 | -0.192 | |
| | Sig. (2-tailed) | 0.503 | 0.84 | 0.884 | 0.304 | 0.5 | 0.846 | 0.409 | 0.312 | 0.393 | 0.086 | 0.971 | 0.132 | 0.112 | 0.368 | |
| | N | 29 | 23 | 25 | 24 | 30 | 24 | 24 | 24 | 24 | 24 | 24 | 24 | 24 | 24 | |
| DNA Lesions Frequency | Correlation Coefficient | -0.13 | 0.044 | -0.031 | 0.219 | 0.128 | -0.042 | -0.177 | 0.216 | 0.183 | 0.357 | 0.008 | 0.317 | 0.333 | 0.192 | -1.000** |
| | Sig. (2-tailed) | 0.503 | 0.84 | 0.884 | 0.304 | 0.5 | 0.846 | 0.409 | 0.312 | 0.393 | 0.086 | 0.971 | 0.132 | 0.112 | 0.368 | <.001 |
| | N | 29 | 23 | 25 | 24 | 30 | 24 | 24 | 24 | 24 | 24 | 24 | 24 | 24 | 24 | 30 |

### Supplementary Table 3: Correlation coefficients and p values of the analysis of the mouse study.

| | | Plasma Ferritin Conc. | Tissue Iron Conc. | Plasma Ferritin Conc. | Tissue Ferritin Conc. | Complex I Activity | Complex II Activity | Complex III Activity | Complex IV Activity | Complex V Expression | Complex III Expression | Complex IV Expression | Complex II Expression | Complex I Expression | mtDNA Copy No. | DNA Lesion Frequency | IRP1 Binding Activity |
|---|---|---|---|---|---|---|---|---|---|---|---|---|---|---|---|---|---|
| Tissue Iron Conc. | Correlation Coefficient | .221* | | | | | | | | | | | | | | | |
| | Sig. (2-tailed) | 0.041 | | | | | | | | | | | | | | | |
| | N | 86 | | | | | | | | | | | | | | | |
| Plasma Ferritin Conc. | Correlation Coefficient | -0.125 | -0.067 | | | | | | | | | | | | | | |
| | Sig. (2-tailed) | 0.247 | 0.538 | | | | | | | | | | | | | | |
| | N | 88 | 86 | | | | | | | | | | | | | | |
| Tissue Ferritin Conc. | Correlation Coefficient | 0.07 | 0.117 | -.331** | | | | | | | | | | | | | |
| | Sig. (2-tailed) | 0.554 | 0.324 | 0.004 | | | | | | | | | | | | | |
| | N | 74 | 73 | 74 | | | | | | | | | | | | | |
| Complex I Activity | Correlation Coefficient | -0.199 | 0.04 | -0.007 | 0.099 | | | | | | | | | | | | |
| | Sig. (2-tailed) | 0.097 | 0.743 | 0.952 | 0.448 | | | | | | | | | | | | |
| | N | 71 | 71 | 71 | 61 | | | | | | | | | | | | |
| Complex II Activity | Correlation Coefficient | -0.076 | -0.23 | -0.011 | -0.193 | -0.03 | | | | | | | | | | | |
| | Sig. (2-tailed) | 0.537 | 0.059 | 0.928 | 0.15 | 0.818 | | | | | | | | | | | |
| | N | 68 | 68 | 68 | 57 | 62 | | | | | | | | | | | |
| Complex III Activity | Correlation Coefficient | -0.049 | -0.068 | .310** | -.615** | -0.053 | -0.166 | | | | | | | | | | |
| | Sig. (2-tailed) | 0.681 | 0.57 | 0.008 | <.001 | 0.674 | 0.19 | | | | | | | | | | |
| | N | 73 | 72 | 73 | 62 | 65 | 64 | | | | | | | | | | |
| Complex IV Activity | Correlation Coefficient | 0.036 | -0.01 | 0.157 | -0.057 | -.279* | .254* | -0.108 | | | | | | | | | |

| | | | | | | | | | | | | | | | |
|---|---|---|---|---|---|---|---|---|---|---|---|---|---|---|---|
| | Sig. (2-tailed) | 0.757 | 0.932 | 0.173 | 0.65 | 0.02 | 0.041 | 0.375 | | | | | | | |
| | N | 77 | 76 | 77 | 65 | 69 | 65 | 70 | | | | | | | |
| Complex V Expression | Correlation Coefficient | -0.154 | -0.191 | .309** | -0.178 | -0.117 | 0.074 | .294* | 0.218 | | | | | | |
| | Sig. (2-tailed) | 0.172 | 0.092 | 0.005 | 0.147 | 0.333 | 0.55 | 0.012 | 0.059 | | | | | | |
| | N | 80 | 79 | 80 | 68 | 70 | 67 | 73 | 76 | | | | | | |
| Complex III Expression | Correlation Coefficient | -0.199 | -0.166 | .385** | -.279* | -0.178 | 0.058 | .317** | 0.097 | .825** | | | | | |
| | Sig. (2-tailed) | 0.077 | 0.143 | <.001 | 0.021 | 0.139 | 0.639 | 0.006 | 0.405 | <.001 | | | | | |
| | N | 80 | 79 | 80 | 68 | 70 | 67 | 73 | 76 | 80 | | | | | |
| Complex IV Expression | Correlation Coefficient | -0.111 | 0.037 | 0.132 | 0.002 | -0.073 | -0.006 | 0.075 | -0.073 | .599** | .680** | | | | |
| | Sig. (2-tailed) | 0.326 | 0.744 | 0.242 | 0.984 | 0.547 | 0.959 | 0.529 | 0.534 | <.001 | <.001 | | | | |
| | N | 80 | 79 | 80 | 68 | 70 | 67 | 73 | 76 | 80 | 80 | | | | |
| Complex II Expression | Correlation Coefficient | 0.097 | -0.092 | 0.152 | -0.19 | -.254* | 0.196 | 0.06 | -0.007 | .394** | .424** | .495** | | | |
| | Sig. (2-tailed) | 0.39 | 0.419 | 0.177 | 0.12 | 0.034 | 0.112 | 0.613 | 0.953 | <.001 | <.001 | <.001 | | | |
| | N | 80 | 79 | 80 | 68 | 70 | 67 | 73 | 76 | 80 | 80 | 80 | | | |
| Complex I Expression | Correlation Coefficient | 0.096 | -0.105 | 0.077 | -0.083 | -.249* | .273* | -0.138 | 0.171 | .393** | .418** | .429** | .814** | | |
| | Sig. (2-tailed) | 0.396 | 0.357 | 0.495 | 0.5 | 0.038 | 0.025 | 0.244 | 0.139 | <.001 | <.001 | <.001 | <.001 | | |
| | N | 80 | 79 | 80 | 68 | 70 | 67 | 73 | 76 | 80 | 80 | 80 | 80 | | |
| mtDNA Copy No. | Correlation Coefficient | -0.108 | -0.085 | -0.044 | -0.129 | 0.144 | -0.04 | 0.025 | -0.202 | -0.08 | -0.085 | -0.063 | -0.112 | -0.004 | |
| | Sig. (2-tailed) | 0.325 | 0.442 | 0.688 | 0.28 | 0.239 | 0.748 | 0.835 | 0.082 | 0.486 | 0.457 | 0.583 | 0.331 | 0.97 | |
| | N | 85 | 84 | 85 | 72 | 69 | 67 | 72 | 75 | 78 | 78 | 78 | 78 | 78 | |
| DNA Lesion frequency | Correlation Coefficient | 0.108 | 0.085 | 0.044 | 0.129 | -0.144 | 0.04 | -0.025 | 0.202 | 0.08 | 0.085 | 0.063 | 0.112 | 0.004 | -1.000** |
| | Sig. (2-tailed) | 0.325 | 0.442 | 0.688 | 0.28 | 0.239 | 0.748 | 0.835 | 0.082 | 0.486 | 0.457 | 0.583 | 0.331 | 0.97 | <.001 |

| | | | | | | | | | | | | | | | | |
|---|---|---|---|---|---|---|---|---|---|---|---|---|---|---|---|---|
| | N | 85 | 84 | 85 | 72 | 69 | 67 | 72 | 75 | 78 | 78 | 78 | 78 | 78 | 85 | | |
| IRP1 Binding Activity | Correlation Coefficient | -0.147 | -0.085 | -0.066 | .333** | -0.049 | 0.085 | -0.186 | -0.011 | 0.038 | 0.02 | 0.088 | -0.009 | 0.057 | -0.115 | 0.115 | |
| | Sig. (2-tailed) | 0.178 | 0.442 | 0.549 | 0.004 | 0.69 | 0.499 | 0.121 | 0.924 | 0.741 | 0.863 | 0.444 | 0.937 | 0.621 | 0.296 | 0.296 | |
| | N | 85 | 84 | 85 | 72 | 69 | 66 | 71 | 75 | 78 | 78 | 78 | 78 | 78 | 84 | 84 | |
| IRP1 Expression | Correlation Coefficient | -0.2 | -0.144 | -0.051 | .312* | 0.233 | 0.156 | -.384** | -0.018 | -0.098 | -0.12 | -0.085 | -0.04 | 0.074 | -0.1 | 0.1 | 0.217 |
| | Sig. (2-tailed) | 0.092 | 0.227 | 0.672 | 0.016 | 0.075 | 0.254 | 0.003 | 0.888 | 0.436 | 0.337 | 0.495 | 0.752 | 0.552 | 0.406 | 0.406 | 0.067 |
| | N | 72 | 72 | 72 | 59 | 59 | 55 | 59 | 66 | 66 | 66 | 66 | 66 | 66 | 71 | 71 | 72 |

STROBE Statement—Checklist of items that should be included in reports of ***cohort studies***

| | Item No | Recommendation | | Page |
|---|---|---|---|---|
| **Title and abstract** | 1 | (*a*) Indicate the study's design with a commonly used term in the title or the abstract | | 1 |
| | | (*b*) Provide in the abstract an informative and balanced summary of what was done and what was found | | 2 |
| **Introduction** | | | | |
| Background/rationale | 2 | Explain the scientific background and rationale for the investigation being reported | | 3 |
| Objectives | 3 | State specific objectives, including any prespecified hypotheses | | 3 |
| **Methods** | | | | |
| Study design | 4 | Present key elements of study design early in the paper | | 3, 4 |
| Setting | 5 | Describe the setting, locations, and relevant dates, including periods of recruitment, exposure, follow-up, and data collection | | 3, 4 |
| Participants | 6 | (*a*) Give the eligibility criteria, and the sources and methods of selection of participants. Describe methods of follow-up | | 3 |
| | | (*b*) For matched studies, give matching criteria and number of exposed and unexposed | | 3, 4 |
| Variables | 7 | Clearly define all outcomes, exposures, predictors, potential confounders, and effect modifiers. Give diagnostic criteria, if applicable | | 5-7 |
| Data sources/ measurement | 8* | For each variable of interest, give sources of data and details of methods of assessment (measurement). Describe comparability of assessment methods if there is more than one group | | 5-7 |
| Bias | 9 | Describe any efforts to address potential sources of bias | | 5 |
| Study size | 10 | Explain how the study size was arrived at | | 4 |
| Quantitative variables | 11 | Explain how quantitative variables were handled in the analyses. If applicable, describe which groupings were chosen and why | | 4 |
| Statistical methods | 12 | (*a*) Describe all statistical methods, including those used to control for confounding | | 7 |
| | | (*b*) Describe any methods used to examine subgroups and interactions | | 7 |
| | | (*c*) Explain how missing data were addressed | | N/A |
| | | (*d*) If applicable, explain how loss to follow-up was addressed | | N/A |

| | | | | |
|---|---|---|---|---|
| | | (*e*) Describe any sensitivity analyses | | N/A |
| **Results** | | | | |
| Participants | 13* | (a) Report numbers of individuals at each stage of study—eg numbers potentially eligible, examined for eligibility, confirmed eligible, included in the study, completing follow-up, and analysed | | 20 |
| | | (b) Give reasons for non-participation at each stage | | 8 |
| | | (c) Consider use of a flow diagram | | 20 |
| Descriptive data | 14* | (a) Give characteristics of study participants (eg demographic, clinical, social) and information on exposures and potential confounders | | Supplementary Table 1 |
| | | (b) Indicate number of participants with missing data for each variable of interest | | 7-8 |
| | | (c) Summarise follow-up time (eg, average and total amount) | | N/A |
| Outcome data | 15* | Report numbers of outcome events or summary measures over time | | 7-8 |
| Main results | 16 | (*a*) Give unadjusted estimates and, if applicable, confounder-adjusted estimates and their precision (eg, 95% confidence interval). Make clear which confounders were adjusted for and why they were included | | Table 1 |
| | | (*b*) Report category boundaries when continuous variables were categorized | | N/A |
| | | (*c*) If relevant, consider translating estimates of relative risk into absolute risk for a meaningful time period | | N/A |
| Other analyses | 17 | Report other analyses done—eg analyses of subgroups and interactions, and sensitivity analyses | | N/A |
| **Discussion** | | | | |
| Key results | 18 | Summarise key results with reference to study objectives | | 9 |
| Limitations | 19 | Discuss limitations of the study, taking into account sources of potential bias or imprecision. Discuss both direction and magnitude of any potential bias | | 10-11 |
| Interpretation | 20 | Give a cautious overall interpretation of results considering objectives, limitations, multiplicity of analyses, results from similar studies, and other relevant evidence | | 11 |
| Generalisability | 21 | Discuss the generalisability (external validity) of the study results | | 9-11 |
| **Other information** | | | | |
| Funding | 22 | Give the source of funding and the role of the funders for the present study and, if applicable, for the original study on which the present article is based | | 12 |

*Give information separately for exposed and unexposed groups.

**Supplementary material 1: The ARRIVE checklist for reporting animal studies**

### The ARRIVE Essential 10

These items are the basic minimum to include in a manuscript. Without this information, readers and reviewers cannot assess the reliability of the findings.

| Item | | Recommendation | Section/line number, or reason for not reporting |
|---|---|---|---|
| **Study design** | 1 | For each experiment, provide brief details of study design including: | |
| | | a. The groups being compared, including control groups. If no control group has been used, the rationale should be stated. | Line 95 - 110 |
| | | b. The experimental unit (e.g. a single animal, litter, or cage of animals). | Line 96 |
| **Sample size** | 2 | a. Specify the exact number of experimental units allocated to each group, and the total number in each experiment. Also indicate the total number of animals used. | Line 95 - 110 |
| | | b. Explain how the sample size was decided. Provide details of any *a priori* sample size calculation, if done. | Line 95 |
| **Inclusion and exclusion criteria** | 3 | a. Describe any criteria used for including and excluding animals (or experimental units) during the experiment, and data points during the analysis. Specify if these criteria were established *a priori*. If no criteria were set, state this explicitly. | Line 96 |
| | | b. For each experimental group, report any animals, experimental units or data points not included in the analysis and explain why. If there were no exclusions, state so. | Line 218 |
| | | c. For each analysis, report the exact value of *n* in each experimental group. | Line 218-222 |
| **Randomisation** | 4 | a. State whether randomisation was used to allocate experimental units to control and treatment groups. If done, provide the method used to generate the randomisation sequence. | Line 100 |
| | | b. Describe the strategy used to minimise potential confounders such as the order of treatments and measurements, or animal/cage location. If confounders were not controlled, state this explicitly. | Line 100-110 |
| **Blinding** | 5 | Describe who was aware of the group allocation at the different stages of the experiment (during the allocation, the conduct of the experiment, the outcome assessment, and the data analysis). | Line 116 |
| **Outcome measures** | 6 | a. Clearly define all outcome measures assessed (e.g. cell death, molecular markers, or behavioural changes). | Line 118 -191 |
| | | b. For hypothesis-testing studies, specify the primary outcome measure, i.e. the outcome measure that was used to determine the sample size. | N/A |
| **Statistical methods** | 7 | a. Provide details of the statistical methods used for each analysis, including software used. | Line 192-201 |
| | | b. Describe any methods used to assess whether the data met the assumptions of the statistical approach, and what was done if the assumptions were not met. | Line 192-201 |
| **Experimental animals** | 8 | a. Provide species-appropriate details of the animals used, including species, strain and substrain, sex, age or developmental stage, and, if relevant, weight. | Line 95 |
| | | b. Provide further relevant information on the provenance of animals, health/immune status, genetic modification status, genotype, and any previous procedures. | Line 95-114 |
| **Experimental procedures** | 9 | For each experimental group, including controls, describe the procedures in enough detail to allow others to replicate them, including: | Line 95-114 |
| | | a. What was done, how it was done and what was used. | |
| | | b. When and how often. | |
| | | c. Where (including detail of any acclimatisation periods). | |
| | | d. Why (provide rationale for procedures). | |
| **Results** | 10 | For each experiment conducted, including independent replications, report: | Table 2 |
| | | a. Summary/descriptive statistics for each experimental group, with a measure of variability where applicable (e.g. mean and SD, or median and range). | |
| | | b. If applicable, the effect size with a confidence interval. | |

### The Recommended Set

These items complement the Essential 10 and add important context to the study. Reporting the items in both sets represents best practice.

| Item | | Recommendation | Section/line number, or reason for not reporting |
|---|---|---|---|
| **Abstract** | 11 | Provide an accurate summary of the research objectives, animal species, strain and sex, key methods, principal findings, and study conclusions. | Line 26-45 |
| **Background** | 12 | a. Include sufficient scientific background to understand the rationale and context for the study, and explain the experimental approach.<br>b. Explain how the animal species and model used address the scientific objectives and, where appropriate, the relevance to human biology. | Line 46-66 |
| **Objectives** | 13 | Clearly describe the research question, research objectives and, where appropriate, specific hypotheses being tested. | Line 59-66 |
| **Ethical statement** | 14 | Provide the name of the ethical review committee or equivalent that has approved the use of animals in this study, and any relevant licence or protocol numbers (if applicable). If ethical approval was not sought or granted, provide a justification. | Line 90-94 |
| **Housing and husbandry** | 15 | Provide details of housing and husbandry conditions, including any environmental enrichment. | Line 59-66 |
| **Animal care and monitoring** | 16 | a. Describe any interventions or steps taken in the experimental protocols to reduce pain, suffering and distress.<br>b. Report any expected or unexpected adverse events.<br>c. Describe the humane endpoints established for the study, the signs that were monitored and the frequency of monitoring. If the study did not have humane endpoints, state this. | N/A<br>N/A<br>Line 110 |
| **Interpretation/ scientific implications** | 17 | a. Interpret the results, taking into account the study objectives and hypotheses, current theory and other relevant studies in the literature.<br>b. Comment on the study limitations including potential sources of bias, limitations of the animal model, and imprecision associated with the results. | Line 216-255<br>Line 310-337 |
| **Generalisability/ translation** | 18 | Comment on whether, and how, the findings of this study are likely to generalise to other species or experimental conditions, including any relevance to human biology (where appropriate). | N/A |
| **Protocol registration** | 19 | Provide a statement indicating whether a protocol (including the research question, key design features, and analysis plan) was prepared before the study, and if and where this protocol was registered. | N/A |
| **Data access** | 20 | Provide a statement describing if and where study data are available. | N/A |
| **Declaration of interests** | 21 | a. Declare any potential conflicts of interest, including financial and non-financial. If none exist, this should be stated.<br>b. List all funding sources (including grant identifier) and the role of the funder(s) in the design, analysis and reporting of the study. | Line 359-361<br>Line 362-365 |
